# The Next Generation Precision Medical Record - A Framework for Integrating Genomes and Wearable Sensors with Medical Records

**DOI:** 10.1101/039651

**Authors:** Daryl Waggott, Anja Bog, Enakshi Singh, Prag Batra, Mark H Wright, Euan Ashley, Dianna Fisk, Anna Shcherbina, Jessica Torresl, Matthew Wheeler, Jason Merker, Carlos D Bustamante

## Abstract

Current medical records are rigid with regards to emerging big biomedical data. Examples of poorly integrated big data that already exist in clinical practice include whole genome sequencing and wearable sensors for real time monitoring. Genome sequencing enables conventional diagnostic interrogation and forms the fundamental baseline for precision health throughout a patient’s lifetime. Mobile sensors enable tailored monitoring regimes for both reducing risk through precision health interventions and acute condition surveillance. In order to address the absence of these data in the Electronic Medical Record (EMR), we worked with the SAP Personalized Medicine team to re-envision a modern medical record with these components. The pilot project used 37 patient families with complex medical records, whole genome sequencing and some level of wearable monitoring. Core functionality included patient timelines with integrated text analytics, personalized genomic curation and wearable alerts. The current phase is being rolled out to over 1500 patients in clinics across the hospital system. While fundamentally research, we believe this proof of principle platform is the first of its kind and represents the future of data driven clinical medicine.

## Background

Electronic Medical Records (EMR) are the core of modern health care systems. Over the last 20 years, they have slowly emerged from paper based records into an essential tool for tracking contact information, patient encounters, quality of care metrics and billing. While they exist as successful tools for administrative purposes, they tend to be technically unsophisticated when it comes to clinical care. Increasingly, high volumes of clinical data such as imaging and genomics are being generated and immediately siloed and often discarded due to lack of central infrastructure. While potentially transformative to clinical care, the demands of stability and curse of legacy leave EMR systems limping at a glacial pace towards cost efficiency and medical innovation.

The complexities of the American health care system were discussed at length in *‘The Anatomy of Health Care in the United States’* (Moses et al. 2013). The system is staggering. As of 2011, 15.7% of the US workforce was employed in health care, with expenditures of $2.7 trillion (17.9% of GDP). Even with increases in resources, prices are rising (hospital charges +4.2%/yr, drugs and devices +4.0%/yr) and basic quality metrics such as life expectancy are behind other developed nations. The authors discuss several major forces of change in health care that speak directly to the challenge and opportunity that this framework works to address. First, consolidation through horizontal aggregation of similar entities has formed vertically integrated heterogeneous and generally insular organizations. Second, information technology investments have focused on administrative efficiencies and improving quality of care. While admirable objectives, these goals have a potential for significant improvement. For example, US billing costs are 13% vs 6.6% in Canada. In order for coordinated clinical care to be realized, information needs to be collected, standardized, shared in a digital format and ultimately innovative clinical applications need to be created. This is not trivial in a large, slow-moving, regulatory-burdened system.

The existing medical record is opaque to most modern analytical tools (Wang et al. 2015; Tamang et al. 2015; Jung et al. 2015; Curé et al. 2015). Examples from first hand experience with multiple large EMR installations include the following: 1) Clinicians struggle to search for their own patients using keywords, and in many cases cannot complete this simple task. Clinical encounters are recorded as a series of *‘notes’* to form a timeline. 2) Notes are free text and subjective to each clinician of varying quality and lacking any type of semantic structure. 3) External reports, images or assays are stored as bitmap data blobs without metadata and no direct method of querying. Even if clinical labs or diagnostic tests have been run, they are often entered manually and therefore riddled with errors or simply missing. Shockingly, extracting the relevant disorder or ICD9 code is in many cases not even possible without manual intervention (Mark and David 2015). However, there have been developments to embed greater support for medical ontologies to better capture diagnosis and symptoms (Girdea et al. 2013; Robinson et al. 2008). There are also a number of disruptively nimble EMR platforms such as the cloud based Practice Fusion and Athena Health (http://www.practicefusion.com/; http://www.athenahealth.com/).

The concept of a linked medical record has been worked on with varying success in most national healthcare systems, but has been difficult within the balkanized nature of the US system (Weber, Mandl, and Kohane 2014). An area that has not been not been tackled sufficiently is the integration of large, complex data within a patient’s actual record. In this document we propose an alternative model focused on deep data centric patient integration. More specifically, we propose a framework for linking patient medical records with genomic data and wearable sensor monitoring. A key dependency for this integration is the ability to transform unstructured clinical encounters into informatically amenable data structures (Liao et al. 2015). This transformation will enable contextual interpretation of genomic results and wearable sensor monitoring for continuous care.

Clinical genetic testing is quickly moving from single marker tests to large multi gene panels digitally captured from exome and whole genome sequencing (Yang et al. 2014; Lee et al. 2014; Zhu et al. 2015). The inclusion of genomics is changing and some specialized systems have been incredible examples of future possibilities (Dumitrescu et al. 2015; Alterovitz et al. 2015). However, most major EMRs have no capacity to store genomic data, and if so, only maintain high level metadata or a few variants that are deemed reportable. Even when genomics is collected, it tends to be stored as a report relating a single key finding to a specific diagnostic test. Several reasons (beyond ethical) for exclusion include informatic practicality (100s of millions of data points per patient) and lack of clinical applicability, utility or genetic actionability. The former is a technical issue that we address in our framework, while the latter is more subtle and discussed below.

First, genomic sequencing typically originates as a specific diagnostic test and therefore belongs solely within the scope of the original clinical purpose. While technically this is logical, it highlights that genome sequencing should be part of patient’s core medical record, shared among clinicians and supplemented or improved over time. Second, clinical actionability is dependent on the diagnostic context and fails to recognize the huge potential for application in multiple health domains over the lifetime of a patient. In addition, clinically relevant genomic information is systematically evolving through initiatives such as ClinGen. The arguments against storing genomic data are somewhat circular. That is, the future clinical value of genomic data is dependent on collection and collection is argued against by its value proposition. Third, the current clinical record by definition is an atomic unit specific to a patient as defined by HIPAA. This can be at odds with the current clinical genetics diagnostic process which depends on inclusion of family members or patients with similar conditions for diagnostic power (Dewey et al. 2011). Moreover, ACMG guidelines on reporting incidental findings recommend notifying family members of potential carrier status, which is not possible using this model. Storing all of a patient’s clinical data within the EMR would therefore entail risk of future incidental findings and consequent ethical and resource obligations. We systematically tackle the practical issues of storing genomes within the EMR in our framework.

Mobile health is in its infancy relative to genomics within clinical practice. The vast majority of devices are consumer oriented with sophisticated sensor collection of health information on activity, sleep patterns and heart rate. With current clinical practice these metrics are only opportunistically captured at discrete time points. This fails to capture latent risk situations such as changes in magnitude or variability over time or under specific conditions. As an example, atrial fibrillation, which is a major risk factor for stroke, has an estimated prevalence of ∼2% and growing rapidly (3.7-4.2% >49yrs) (Colilla et al. 2013; Chugh et al. 2014; Zoni-Berisso et al. 2014). The presence of the condition is unknown to most patients due to gradual onset or infrequent episodes. For those that do have a moderate or early stage diagnosis, very little is done to continuously monitor and consequently prevent severe events. Similarly, cardiomyopathy, which affects 0.2% of the population, is a major risk factor for AF and sudden death, but is rarely monitored outside of clinic. Wearable sensors are capable of making these measurements and providing affordable non-invasive monitoring of patients by clinicians. While many clinics are working on innovative applications for these data, they struggle with not being able to incorporate the results into the clinical record in a meaningful integrated way. For example, patients with elevated risk could be monitored more closely and targeted for aggressive engagement strategies. A modern clinical record is crucial for this type of intelligent profiling *i.e.* integrating genetic risk predisposition, observed subclinical risk factors such as stationary behavior, presence of comorbidities or taking a medication with predicted adverse events based on a specific genotype.

In this document we describe an analytical framework for integrating classical EMRs with genomics and wearables to create the next generation clinical record. All of this can be done without replacing the underlying EMR investment. We believe the future of healthcare is going to be heavily dependent on granular patient measurement with continuous monitoring and an emphasis on machine learning based algorithms to inform precision health. The solution is to re-imagine what an integrated system would look like with these core fundamentals built in. To the best of our knowledge this is the first application of integrating genomics and wearable sensor data with medical records for actual patients.

## Results and Features

### Patient Triage

A key challenge to a system that integrates many users, patients and data is the act of task or key information prioritization. Upon logging into the framework a clinician or informatician will be presented with patients ranked by two main criteria. First, patients will be ranked relative to the number of outstanding tasks. Tasks could include reflex bioinformatic pipelines, various aspects of variant/gene curation, Sanger validation or additional test ordering. Second, patients will be ordered relative to key information or alerts pertinent to their condition or general health. The representation of the alerts and tasks will be specific to the roles and assigned patients for each system user. An example screenshot can be seen in Figure 1.

**Figure 1.**
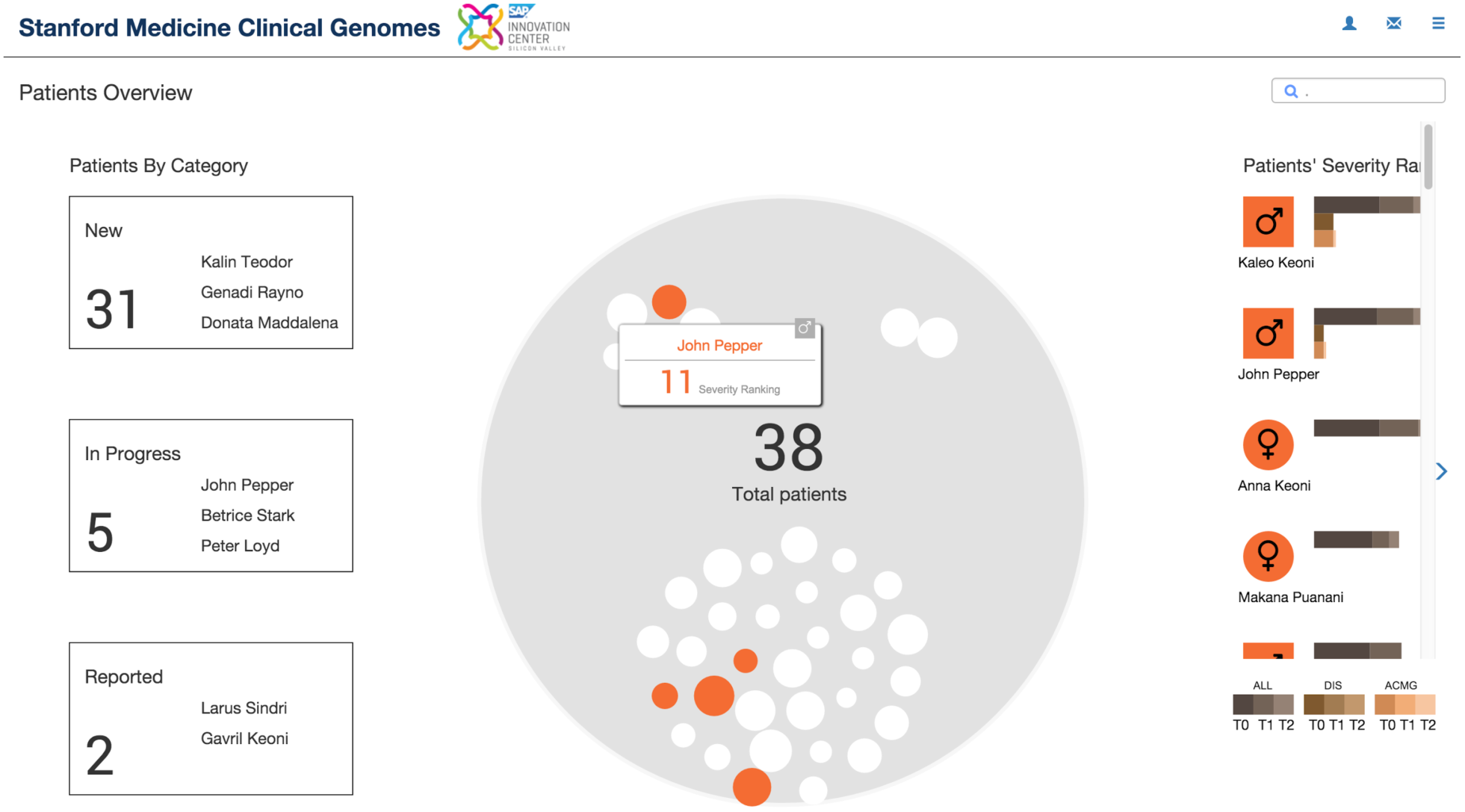
Patient Triage

### Patient Dashboard

The patient triage process then directs a user to a patient dashboard. This is a simplified yet rich overview of a patient’s integrated precision medical record. Key elements include an interactive patient clinical timeline, conventional lab results, wearable sensor alerts and genomic summaries. The genomic summaries are stratified into key clinical findings (either incidental, actionable or condition specific), drug response variants, ABO/HLA genotypes and global ancestry. Each of the elements can be selected for more in-depth review and will be discussed below. An example screenshot can be seen in Figure 2.

**Figure 2.**
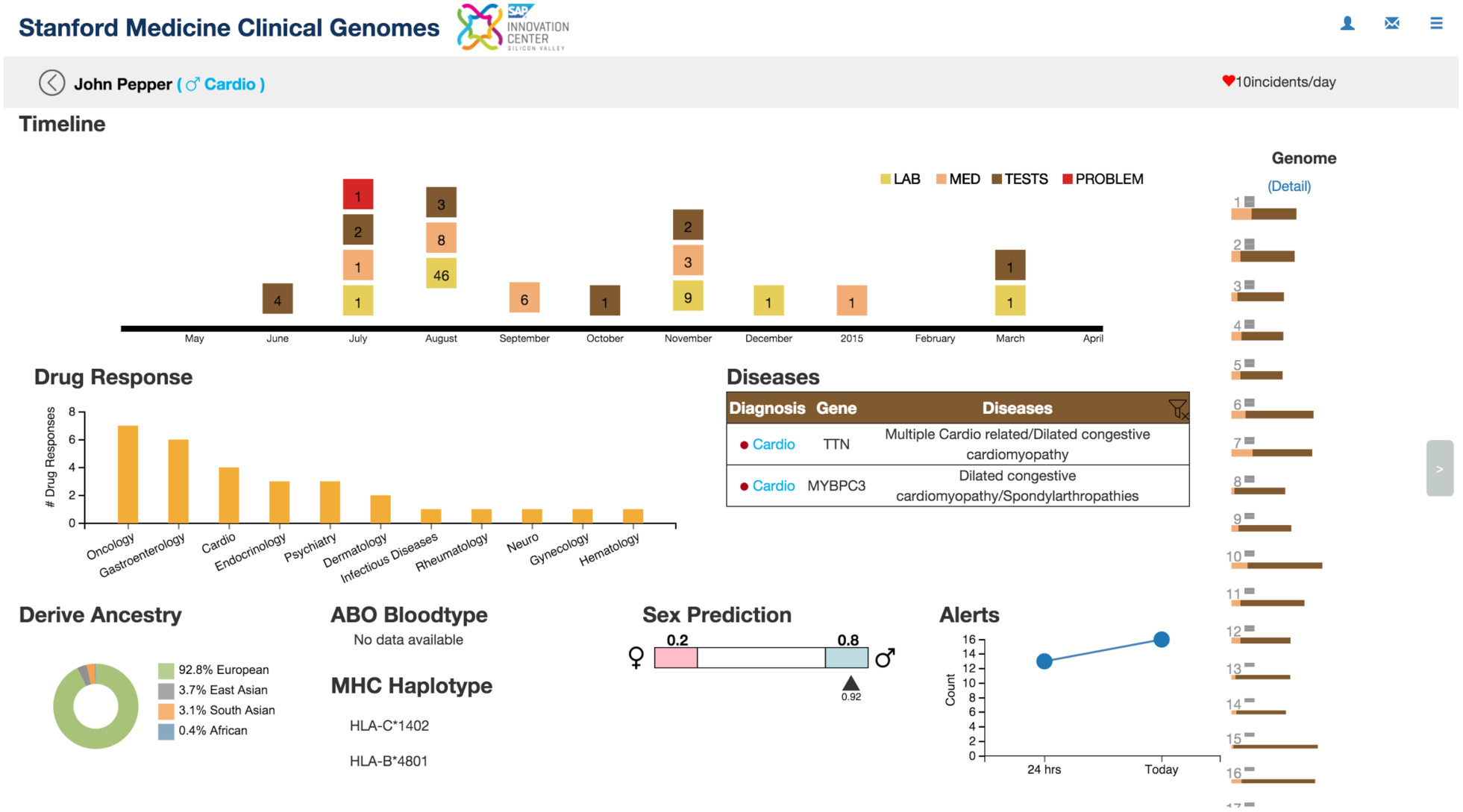
- Patient Dashboard

### Scalable Genome Storage

First and foremost, the infrastructure was designed to scale to handle all genomes for all patients within a large national healthcare system. With SAP HANA as the underlying database platform, this system can scale to store genomic data from thousands or even millions of patients.

Complex annotation joins and aggregate queries are on the order of seconds. Initial loading of genomic data from VCF files currently takes approximately 5 ½ minutes per whole genome (∼10M variants). This step happens only once. It is anticipated that we can dramatically improve these load times by not parsing and temporarily storing all the INFO and FORMAT tags present in the VCF. Ideally, a variant calling pipeline and corresponding output format is specifically optimized for downstream storage. In this scenario, only essential genotype information would be generated from the pipeline with the assumption that annotation and quality metrics can be cheaply calculated on the fly. The genome data model can be reviewed in more detail in supplementary file 1.

We have not currently benchmarked our platform against other early stage projects designed to scale genomic storage to thousands and even millions of genomes. Several examples of these projects include <u>GLnexus</u>, <u>GQT</u> and <u>BGT</u>. <u>GLnexus</u> is a research project from DNAnexus which emphasizes incremental n-of-1 storing of genomes as likelihoods for optimized batch calling. <u>BGT</u> developed by Heng Li is a promising format which aims to be more compact in size, more efficient to process, and more flexible on query than conventional BCF2 format. <u>GQT</u> developed by Aaron Quinlan and Ryan Layer has been described in a recent Nature Methods paper <u>(Layer</u> <u>et al. 2016)</u>. Notably, there were substantial performance gains relative to conventional BCF2 and PLINK2 formats (443-fold in some cases).

### Automated Genetic Curation

Automated genetic curation of risk features and pathogenic variants is central to our vision of the precision medical record. In this framework the medical record is mined for key terms which inform candidate gene lists for variant prioritization. The objective is to empower curation experts and clinicians to focus on high value tasks and not data wrangling.

The heart of the automated curation is based on software we developed called *Sequence To Medical Phenotypes* (STMP) (Dewey et al. 2015). The core module tiers variants within a geneset according to potential pathogenicity. The primary tiers are (1) clinvar pathogenic 2+ stars (2) loss-of-function (LOF) or canonical splice (3) High across species amino acid conservation site (4) Predicted protein deleterious across multiple algorithms. Additional STMP modules focus on prioritization-based segregation within a pedigree and pharmacogenomic (PGx) haplotyping using a Hidden Markov Model (HMM). Preloaded gene lists exist for numerous conditions and classes of actionability. Using STMP within the SAP HANA system allows for substantially improved query performance and simplified management of the large number of external database dependencies. Within the patient dashboard of Patient X (Figure 2), note that the identified disease, candidate gene and variant from the automated prioritization agreed with previous manual curation efforts (Cardiomyopathy: *MYBPC3*).

As the underlying medical record changes and genetic annotation evolves, so too does the curation. Most importantly, the genomic information is directly in the medical record whereby relevant information is “pushed” to the clinician and not manually “pulled” by a curator. The concept of continuous annotation updates and “push” curation, makes re-analysis an automated ongoing task. Figure 3 shows an example of tiered variant prioritization along with a query interface for adjusting thresholds *i.e.* allele frequency, inheritance model and gene tolerance.

**Figure 3.**
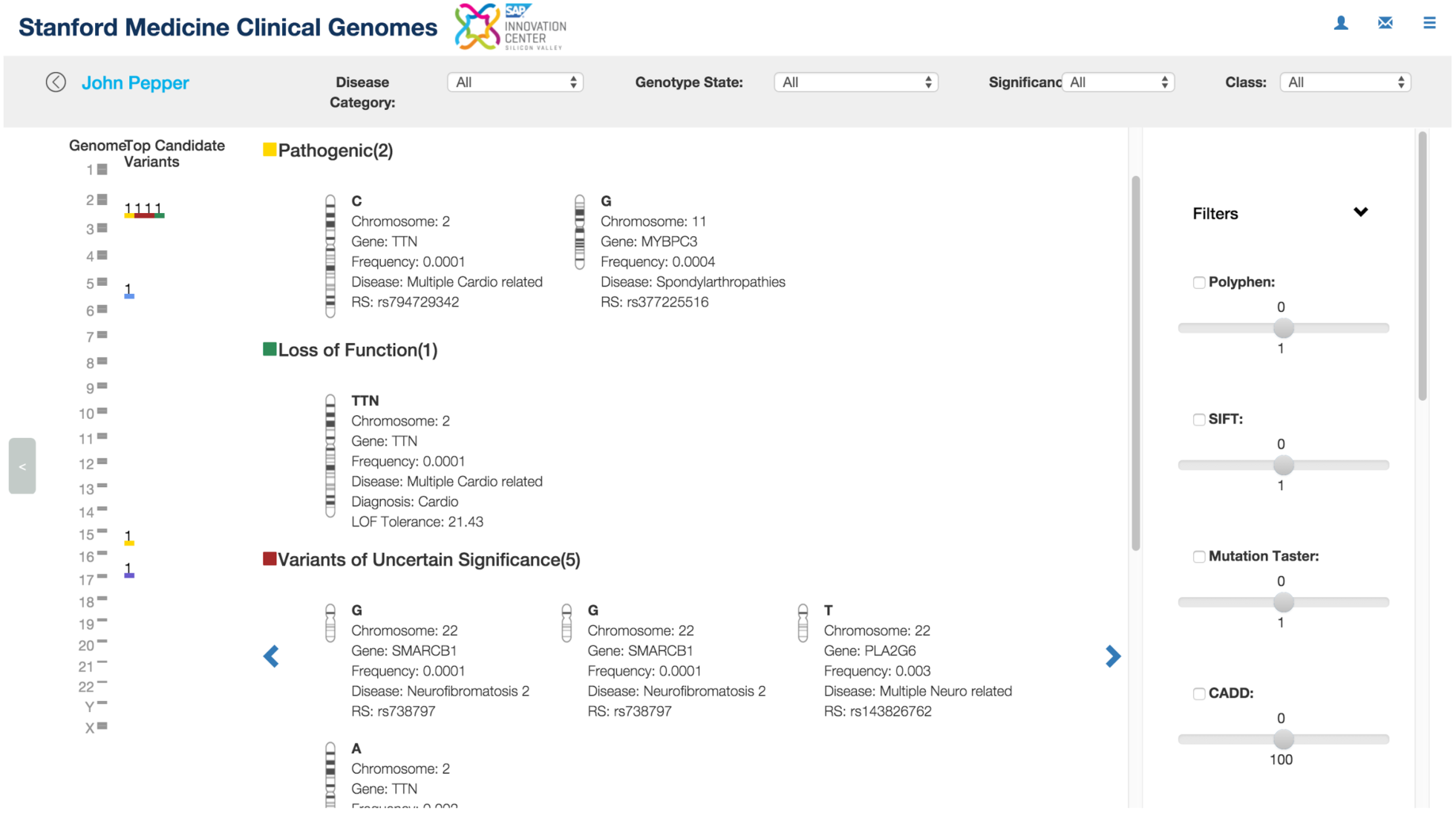
- Variant and Gene Curation

Upon automated prioritization candidate genes and their respective variants can be reviewed independently with extensive opportunity for comments/citations to improve ongoing knowledge base development. Variants can be queried for frequency and clinical associations across all other patients in the system. In fact, these aggregate queries can be included in clinical genetics reporting. Templates exist not only for final clinical genetic reports but intermediate presentations for team discussion. All reports contribute back to the knowledge base for improved future curation.

Beyond complexities around billing, the primary limitation to scaling genomic services in a clinical setting is time spent on curation of genes and variants. The personnel costs alone are substantial (1 genetic counselor / biocurator per case per week). Our approach of integrating genomics into the medical record with substantial back end analytics simplifies human resource burden, streamlines the process and ultimately reduces cost.

### Complex Genomic Annotation

The genome provides a large amount of accessible information relevant to clinical care. Single diagnostic tests for pathogenic variants ignore the use of this rich information for ongoing precision health. As such, we added a number of annotations that require combining information across multiple variants. Several examples include: ABO/HLA haplotypes, PGx haplotypes, sex, relatedness and ancestry. Figure 4 shows a summary of how variants can be interpreted in the context of family segregation and local ancestry (Maples et al. 2013). Additionally, genetic risk scores for common disease are in the process of being added.

**Figure 4.**
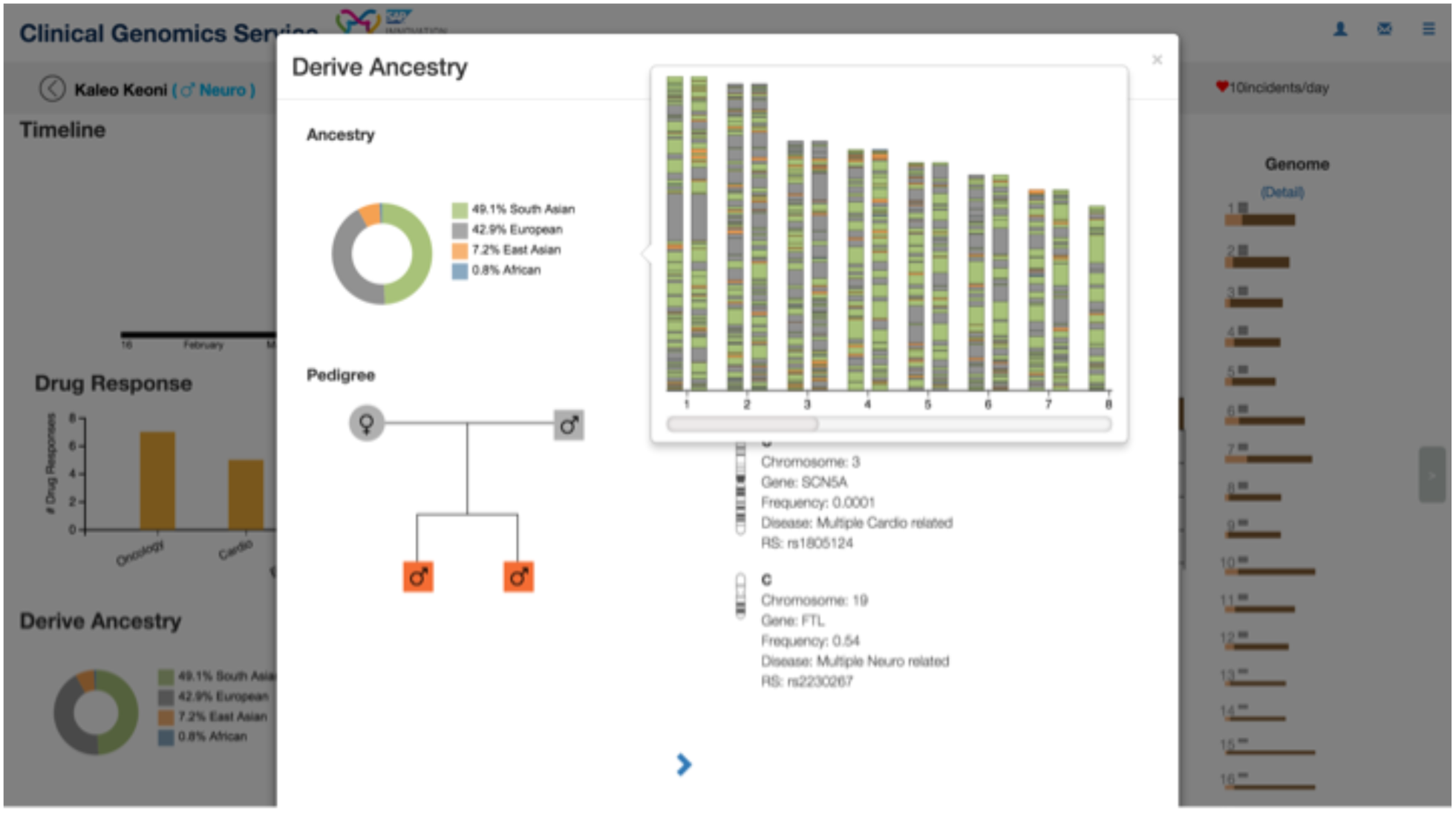
Complex Genomic Annotation

### Real Time Wearable Sensor Monitoring

Figure 5 shows a two day window of Patient X monitored using an Apple watch (3118 observations). Patient X suffered from cardiomyopathy, and therefore any heart rate abnormalities would be of clinical interest. We chose to focus on defining a specific metric for HR relative to predicted energy expenditure. If the patient deviates from the expected relationship an alert is triggered and this is relayed back to the medical record and monitoring physician. You’ll notice that toward the right of the figure there exists an increased density of elevated HR relative to predicted energy expenditure. This reached our threshold of setting off an alert. Several caveats include: (1) Our data was collected with up to a 48 hour synchronization lag. (2) Based on a previous clinical validation we found Apple HR reasonably accurate, but energy expenditure (especially older algorithms based solely on tri-axial accelerometer sensors) was poor *(Consumer Wearable Device Validation with Clinical Standards - in preparation)*. We are currently working on a core set of personalized sensor metrics that could be applied to different health/disease states.

**Figure 5.**
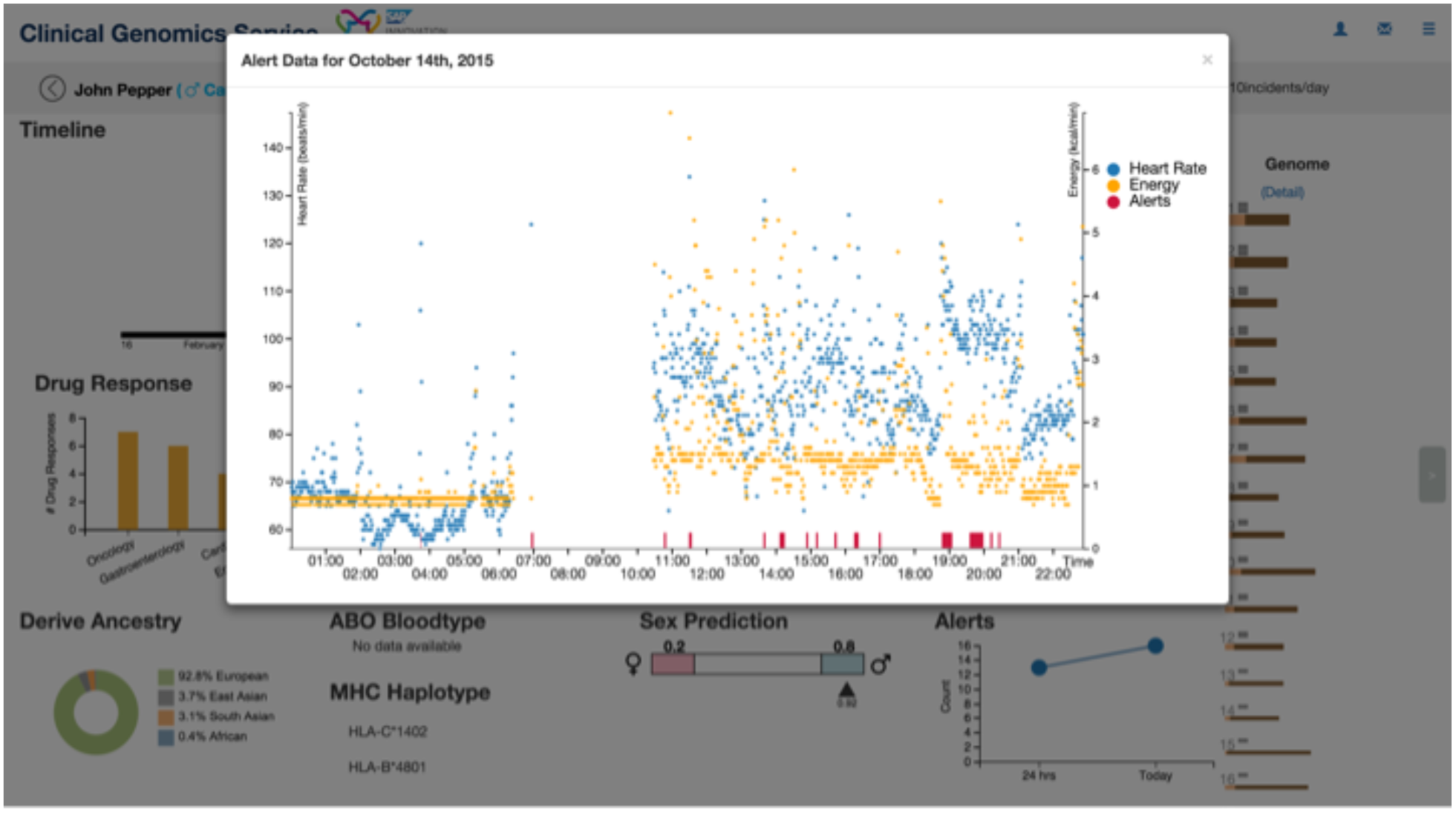
HR and Energy Expenditure Monitoring

### EMR Adapters for Augmentation not Replacement

Stanford Medicine has two independent installations of EPIC. Access to the records are typically restricted to limited forms with no bulk import/export. Our initial pilot of this framework focused on a static EPIC export using the STRIDE platform (Lowe et al. 2009). This approach is only suitable for research and not production. For clinical utility, real time bulk access needs to be enabled. SAP has partnered with the American Society of Clinical Oncology (ASCO) to develop CancerLinQ, a cloud-based big data platform. (<u>Sapphire_2015</u>; <u>ASCO_Connection</u>; <u>ASCO_Meeting_2016</u>). CancerLinQ will enable the secure analyses of patient data from millions of electronic health records from all participating clinics. The first version of CancerLinQ will represent data from approximately 500,00 patients across 15 vanguard practises. To enable CancerLinQ, SAP is developing adapters to extract, transform & load data from all the different EMR systems and harmonize into one single system. These adapters cover the majority of major EMR platforms and are strictly compliant with HIPAA/PHI regulations. As such, data flow is primarily pulled from the EMR and not pushed back. One highly valuable byproduct of the use of abstraction is that unstructured EMR data *i.e.* clinical notes, is transformed into structured information such as key terms and timelines. The use of EMR adapters serves to augment without changing the underlying informatic health structure such as billing and operations.

### Text Mining of Disease Terms

The majority of a clinical encounter is recorded as a ‘clinical note’. This is primarily free text with a large amount of heterogeneity in quality/quantity between clinicians. In a significant number of cases we reviewed, there was no clear or easily machine accessible differential diagnosis or structured clinical traits/symptoms. This information therefore required extraction from the clinical notes. Based on SAP HANA’s natural language processing capabilities, we built a natural language processing pipeline to extract several key clinical features (“MRI Demo,” n.d.; “MRI Accelerate Personalized Trx” 2014). The primary objective was to extract candidate diagnosis. For the most part this depended on identifying ICD9 codes and disease/diagnostic terms *i.e.* MESH values such as cardiomyopathy. The terms were then assigned to a timeline where they were mapped to candidate gene sets using tools such as DisGenNET, OMIM and ClinVar. When possible, well curated disease gene panels such as GeneDx or Invitae would be selected. If a candidate gene panel was not available an *in silico* panel would be created based on the described ontology lookup. The digital candidate gene panel is incredibly important for specificity during curation.

The disease terms further act to tune the use of wearable sensor data collection. Different patient classes would have different algorithms and relevant clinical alerts. For example, a cardiology patient will have a number of HR and exertion measurements whereas a neurology patient may have more of a focus on motility, fine-motor control and coordination.

### Patient Timeline and the Longitudinal Genome

A concept that is not fully developed in our EPIC medical record installation is the patient timeline. A comprehensive overview of each individual patient’s medical history was displayed in a graphical timeline, making it easy to access each patient interaction. The longitudinal perspective of patient encounters and treatment is important for better organization, capturing change in health over time, integration with sensor monitoring and maintenance of clinical care. From the genomics standpoint, automated genetic curation is relevant to a specific window of time. As expected the relevance of different genomic features will change over time within a patient. For example cardiovascular risk or treatment vs oncology variants would be used independently in time and by practitioner. Improved focus on a longitudinal view of the patient’s health helps emphasize the relevance of prospective health as opposed to reactionary treatment. Moreover, it demonstrates that the utility of genomics is multifaceted and should exist as a fundamental baseline within all medical records. Finally, continuous sensor monitoring can be adapted over time and simply correlated with clinical management. Fundamentally, patient health is not cross sectional and we show a proof of principle for the utility of its incorporation into the medical record.

### Version-ome

The fundamental algorithms and supporting annotation used for calling and interpreting features is evolving at terrific pace. In addition, over a patient’s lifetime it is possible to have multiple tissues sequenced for different purposes such as germline risk or tumor profiling. Given that a person’s genomic profile is not static, we chose to explore the concept of a “version-ome”. Essentially, can we apply conventional database management techniques to allow for regulatory-compliant, evolving representation of one’s genome. For example, access date stamps, storing incremental changes on updates and strictly versioning the hundreds of secondary data sources used for curation. This versioning allows for reproducible curation queries, so that specific variants can be revisited years in the future. For example, any variant found in a clinical report will have have an auditable query associated with it.

### Contextual Clinician Alerts

Another key design thought for our next generation medical record system is some degree of automated intelligence. With semi-automated prioritization/curation the system can alert clinicians to only the most fundamental changes in information relevant to a patient’s current condition or latent risk. For example, cardiovascular labs become elevated and patient’s genomic risk is high. Subsequently, PGx variants both in terms of efficacy/safety could inform treatment choice. Also, as underlying ClinVar pathogenicity assertions evolve, an alert could be initiated if a diagnostically relevant threshold is met for a VUS.

Similarly, alerts can be generated based on wearable sensor data streams. As patients move outside of personalized tolerance zones (based on EMR tuning), an alert to a clinician would be initiated. More systematic trends could be summarized during regular clinical encounters and would not trigger alerts. In or proof of concept development we focused on alerts examples relative to clinical genomic insight and how that could inform wearable monitoring. Expanding focus to analysis of already existing medical data streams offers substantial opportunity but was out of scope (Bailey et al. 2013; Mao et al. 2012).

## Methods and Technology

### Development Process

The process of joint development between SAP Personalized Medicine and Stanford Medicine followed design thinking methodologies (Brown 2009; “This Is Design Thinking,” n.d.). First, the SAP team conducted expert interviews with stakeholders at Stanford involved in genome research and clinical care, including genetic counselors, researchers, statisticians and physicians. Key challenges were explored and presented back to the Stanford team. A direction and scope were agreed upon and then development began following scrum agile methodologies (Sutherland and Sutherland 2014). During development, the prototype was presented back to the Stanford stakeholders which contributed to several new iterations of the features within the prototype.

### Cohort

During development three initial cohorts of subjects were tested. Initial prototyping was done on 1000 genomes data (2,504 WGS), Personal Genome Project and 250 exomes from the ELITE project (elite.stanford.edu). The above samples were not patients and lacked accessible electronic medical records and wearable data. Subsequently, we focused development on a cohort of 37 whole genomes from a Stanford Medicine clinical genome pilot project and 8 from the Stanford Undiagnosed Disease Network clinical site. All of these patients had Stanford medical records which were accessed (de-identified) under a combination of 4 IRB protocols. Patients were a combination of cardiology, neurology, inherited cancer and multi-system presentations. A subset of the patients were selected for continuous mHealth sensor monitoring using three consumer devices (apple watch, basis peak and polar with chest strap).

### EMR

The existing medical record system at Stanford Medicine is EPIC and a technology called STRIDE was used to map table structure and export data (Lowe et al. 2009). Among the 37 pilot patients there were 124 detailed diagnosis, >1400 medications, >22,000 lab results and >1000 procedures.

### Backend Architecture

The data model, analytical engine and user interface were all implemented using the SAP HANA development platform, a flexible, multi-purpose in-memory data management and application platform. By providing advanced capabilities, such as text analytics, spatial processing and data virtualization on the same architecture, it further simplifies application development and processing across big-data sources and structures. The setup consisted of a distributed 10 node SAP HANA landscape (9 active and one hot standby). Each node was outfitted with 1TB of memory and 80 Intel(R) Xeon(R) CPU E7 - 4870 @ 2.40GHz. More specific details on performance metrics for in memory systems can be found in the following references.

*The Impact of Columnar In-Memory Databases on Enterprise Systems (Plattner 2014) SAP HANA Database - Data Management for Modern Business Applications (Färber et al. 2012)*

*A Course in In-Memory Data Management (Plattner 2013)*

### Genetic Data Model

All genetic data was extracted from variant call format files (VCF) and transformed into a relational data model within SAP HANA. The VCF file format is incredibly extensible in order to allow tool specific annotation and QC metrics (“[PDF]The Variant Call Format (VCF) Version 4.2…- Samtools,” n.d.). This flexibility creates a large amount of heterogeneity in data parsing and overall representation. Moreover, multiple variants or genotypes can exist at a single position, especially within large cohorts. Additionally, any one subject has 2 copies of DNA or alleles at all 3 billion positions. All of these positions potentially could have annotation and QC data. Finally, for the system to be scalable it needs to be able to handle 100s of thousands or even millions of genomes. Together, these challenges emphasize the need for a robust yet streamlined data model.

To address these challenges we focused on storing the genotypes with only the most fundamental QC metrics. Our assumption was that a subset of QC metrics could be generated on the fly as requested. Conventional annotations (i.e. ClinVar, protein domain, eQTL site…) were stored as tertiary tables for real time querying in order to allow asynchronous updates. The real time nature of queries enabled the dynamic integration of hundreds of annotation sources.

For functional effect prediction (such as performed by Annovar, VEP and SNPEff) we selected a hybrid precompute versus on demand compute model. First, we precomputed all high value SNP positions such as coding, <u>dbSNP</u>, <u>ClinVar</u> and <u>COSMIC</u>. This includes observed and unobserved genotypes in regions of functional importance. Similarly, we calculated all previously observed INDELs. For unobserved INDELs that could not be precalculated, we annotated with transcript, size and frame. As new SNP and INDEL positions are precomputed, they are exported, and loaded for future analyses.

The specifics of the data model can be found in Supplementary File 1 and in the following patent (Patent# 20150234870 - DYNAMIC MAPPING OF EXTENSIBLE DATASETS TO RELATIONAL DATABASE SCHEMAS).

### Genomics Data

**Pipeline**. The genotypes were called using GRCH37-GATK 2.7 best practices.

**Pathogenicity.** Variants were annotated with complete clinvar dumps, including assertions and reviewing organization. Prioritization of pathogenic variants was limited to two plus star likely pathogenic and pathogenic variants. Contradictory assertions and variants of unknown significance were only considered for curation purposes.

Actionable. Variants were determined to be actionable if they were annotated as likely deleterious and in ACMG guidelines or curated lists such as 114 genes proposed by (Dorschner et al. 2013).

**PGx.** <u>FDA labels</u>, <u>PharmaADME</u>, <u>The Human Protein Atlas</u>, CYP alleles

**Risk scores.** Varimed and other validated scores such as Knowles et al. b Sex prediction was based on homozygozity of the X chromosome.

**HLA.** Haplotypes were estimated using several tools based on panels of phased polymorphic SNPs

**Local Ancestry**. Haplotypes were generated using high quality variants present on the Illumina Omni2.5 and phased using ShapeIT with 1000 genomes v3 reference panel. Local ancestry was estimated using a second generation optimization of the RFMIX algorithm (Maples et al. 2013). Briefly, the input phased haplotypes were corrected for statistically inconsistent ancestral and long range LD patterns. Next, a random forest was used to classify the most likely population for each allele within a sliding window. The reference populations were chosen for purity from recent admixture. Global ancestry was assessed by summing all segment assignments and subsequently compared to an extension of ADMIXTURE (Alexander, Novembre, and Lange 2009). The extension is based on projection of a reference population onto the target population of interest. The approach enables robust application to target datasets that contain confounding structure such as relatedness.

### Mobile Data

Three devices were used for data collection: the Apple watch, Basis Peak, and Polar watch with chest strap. Devices were put in collection mode during any modest to strenuous activity. Data was downloaded using device specific APIs. This approach is in the process of being automated with a clinical porting of the app MyHeartCounts (“MyHeartCounts,” n.d.). MHC has enrolled 50,000 users, with the collection of conventional cardiovascular risk factors in combination with mobile based sensor data through HealthKit. Porting a mirror of the back end to Stanford servers will allow for maintenance of regulatory compliance when dealing with patient data. We foresee health care providers managing suites of medical apps that can be prescribed for seamless integration with medical record systems. MHC-clinical is a working example of this model.

## Supporting information

s1-vcf_specification

## Acknowledgments

SAP Labs, LLC including: Anil Ankisettipalli, Surbhi Sheth-Shah, Hameesh Manadath, Jane Luo, Peter Weigt, Janos Brauer General Manager for the Personalized Medicine unit at SAP: Werner Eberhardt

