## Supplementary material for "The Next Generation Precision Medical Record - A Framework for Integrating Genomes and Wearable Sensors with Medical Records": s1-vcf_specification

### VCF Representation in the Database

This document is intended to help template designers understand the mapping of the variant call format (VCF) data to a relational data schema to ultimately write new analysis templates. Structure of the document:

#### Basics of VCF 4.2

VCF is a text-based format to store genetic variant data. The following mapping is based on VCF 4.2[1] and may be subject to change in the future. Listing 1 shows an example for the contents of a VCF file. The file is composed of lines of three formats: (1) metadata lines, (2) one header line, and (3) content lines. The metadata lines start with “##” and they contain information that is used to interpret the contents that follows (lines 1 – 18 in Listing 1). The header line (line 19) starting with “#CHROM” provides the structure of all the following lines, which are the content lines (lines 20 – 24).

```
1    ##fileformat=VCFv4.2
2    ##fileDate=20090805
3    ##source=myImputationProgramV3.1
4    ##reference=file:///seq/references/1000GenomesPilot-NCBI36.fasta
5    ##contig=<ID=20,length=62435964,assembly=B36,md5=f126cdf8a6e0c7f379d618ff66beb2da,
     species="Homo sapiens",taxonomy=x>
6    ##phasing=partial
7    ##INFO=<ID=NS,Number=1,Type=Integer,Description="Number of Samples With Data">
8    ##INFO=<ID=DP,Number=1,Type=Integer,Description="Total Depth">
9    ##INFO=<ID=AF,Number=A,Type=Float,Description="Allele Frequency">
10   ##INFO=<ID=AA,Number=1,Type=String,Description="Ancestral Allele">
11   ##INFO=<ID=DB,Number=0,Type=Flag,Description="dbSNP membership, build 129">
12   ##INFO=<ID=H2,Number=0,Type=Flag,Description="HapMap2 membership">
13   ##FILTER=<ID=q10,Description="Quality below 10">
14   ##FILTER=<ID=s50,Description="Less than 50% of samples have data">
15   ##FORMAT=<ID=GT,Number=1,Type=String,Description="Genotype">
16   ##FORMAT=<ID=GQ,Number=1,Type=Integer,Description="Genotype Quality">
17   ##FORMAT=<ID=DP,Number=1,Type=Integer,Description="Read Depth">
18   ##FORMAT=<ID=HQ,Number=2,Type=Integer,Description="Haplotype Quality">
19   #CHROM POS     ID        REF    ALT     QUAL FILTER INFO
     FORMAT      NA00001        NA00002        NA00003
20   20     14370   rs6054257 G      A       29   PASS   NS=3;DP=14;AF=0.5;DB;H2
     GT:GQ:DP:HQ 0|0:48:1:51,51 1|0:48:8:51,51 1/1:43:5:.,.
21   20     17330   .         T      A       3    q10    NS=3;DP=11;AF=0.017
     GT:GQ:DP:HQ 0|0:49:3:58,50 0|1:3:5:65,3   0/0:41:3
22   20     1110696 rs6040355 A      G,T     67   PASS
     NS=2;DP=10;AF=0.333,0.667;AA=T;DB GT:GQ:DP:HQ 1|2:21:6:23,27 2|1:2:0:18,2   2/2:35:4
23   20     1230237 .         T      .       47   PASS   NS=3;DP=13;AA=T
     GT:GQ:DP:HQ 0|0:54:7:56,60 0|0:48:4:51,51 0/0:61:2
```

[1] 1000 Genomes. VCF 4.2. URL: http://www.1000genomes.org/wiki/analysis/variant-call-format/vcf-variant-call-format-version-42 (last accessed: Jan 24, 2014)

```
24   20      1234567 microsat1 GTC     G,GTCT  50    PASS   NS=3;DP=9;AA=G
     GT:GQ:DP    0/1:35:4       0/2:17:2       1/1:40:3
```
**Listing 1: Example data for VCF 4.2[1] (with added line numbers in front)**

Only a limited subset of the information in the metadata lines is fixed. Most of the information given there is optional and the creator of a VCF file can specify their contents freely following a given basic format.

#### Mapping Description

Since VCF is an "open" format, i.e. users can specify their own fields to store additional data, the mapping needs to accommodate flexibility. The mapping described in the following relies on the ability of a column-oriented database to add new columns to an existing table on the fly during runtime. Thus, the relational database schema evolves over time.

General columns that occur in all tables are the "DATA SET" and "ETL LOAD ID" columns. These ensure the link between imported data and the original VCF files.

##### Metadata Tables

Metadata from VCF files is stored in eight tables: Some of the metadata tables follow a key/value store format instead of the traditional relational model of mapping data to named columns. The rationale behind this is that metadata as mentioned above can be freely specified by VCF file creators and may strongly over a set of files. Yet, the amount of metadata provided is very limited per file. Thus, these tables will grow slowly even using the key/value approach. Furthermore, we assume that the metadata there is rarely used at query time and even if it is being used, it is not necessary to reconstruct the former lines, as they are present in the original VCF file. Such a reconstruction would entail self-joins of tables that would quickly become expensive with growing table sizes.

All lines of a format similar to `##fileformat=VCFv4.2`, that is simple key/value pairs, are mapped to the table "VARIANT SOURCE MASTER". These are for example the lines 1 – 4, and 6 in Listing 1. The lines starting with "##INFO" are mapped to the table "VARIANT INFO MASTER", e.g. lines 7 – 12 in Listing 1, filling the columns according to the keys and values given in the section between the symbols "<",">". All lines starting with "##SAMPLE" are stored in the "VARIANT SAMPLE MASTER" table in the same approach as in the "VARIANT INFO MASTER" table. Other tables following that approach are "VARIANT FORMAT MASTER" (lines starting with "##FORMAT", e.g. lines 15 – 18 in Listing 1), "VARIANT FILTER MASTER" (lines starting with "##FILTER", e.g. lines 13 and 14), and "VARIANT ALT MASTER".

Lines similar to the format of line 5 in Listing 1 are mapped to the table "VARIANT CONTIG MASTER". This table does not have columns for all the possible keys that might occur like the "VARIANT INFO MASTER" table. Instead, all key/value pairs except "ID", are stored in the key/value format. "ID" is stored in the "CONTIG ID" column. The mapping to the table "VARIANT PEDIGREES MASTER" follows the same approach with all lines starting with "##PEDIGREE".

Please see Listing 2 for the names and columns of the metadata tables.

---

**VARIANT_SOURCE_MASTER**
- DATA_SET (string)
- ETL_LOAD_ID (integer),
- ATTRIBUTE_VALUE (string),
- ATTRIBUTE_NAME (string),

**VARIANT_SAMPLE_MASTER**
- DATA_SET (string), ETL_LOAD_ID (integer)
- SAMPLE_ID (string),
- GENOME (string),
- MIXTURE (string),
- DESCRIPTION (string)

**VARIANT_PEDIGREES_MASTER**
- DATA_SET (string), ETL_LOAD_ID (integer)
- NAME (string)
- VALUE (string)

**VARIANT_INFO_MASTER**
- DATA_SET (string), ETL_LOAD_ID (integer)
- DATA_TYPE (string)
- INFO_ID (string)
- NUMBER (string)
- DESCRIPTION (string)

**VARIANT_FORMAT_MASTER**
- DATA_SET (string), ETL_LOAD_ID (integer)
- DESCRIPTION (string),
- DATA_TYPE (string),
- FORMAT_ID (string),
- NUMBER (string),

**VARIANT_FILTER_MASTER**
- DATA_SET (string), ETL_LOAD_ID (integer)
- DESCRIPTION (string),
- FILTER_ID (string)

**VARIANT_CONTIG_MASTER**
- DATA_SET (string), ETL_LOAD_ID (integer)
- CONTIG_ID (string),
- NAME (string),
- VALUE (string),

**VARIANT_ALT_MASTER**
- DATA_SET (string), ETL_LOAD_ID (integer)
- ESCRIPTION (string)
- ALT_ID (string)

---

**Listing 2: Metadata tables**

#### Content Data Mapping

Data from the content lines is recorded in six tables. In addition to the "DATA SET" and "ETL LOAD ID" columns, all these tables store the chromosome "CHROM" and position "POS" where a specific variant has been encountered. A single VCF file will have the same unique combination of "DATA SET" and "ETL LOAD ID" in all its entries in the tables for identification. Since the combination of chromosome and position is not unique in a VCF file, that is, one position in a chromosome can occur in multiple lines in VCF, an extra column called "RECORD COUNT" has been introduced. This column has to be used when joining across tables within VCFs to exactly reproduce the original lines from the VCF file.

The first four tables ("VARIANT MASTER", "VARIANT ALTERNATE", "VARIANT FILTER", "VARIANT ANNOTATION") are of static structure; the last two ("VARIANT INFO" and "VARIANT SAMPLE") have a dynamic structure. These two tables evolve over time in such a way that columns are added when more and more VCF files are loaded into the system that contain their own metadata.

---

**VARIANT_ALTERNATE**

- DATA_SET (string), ETL_LOAD_ID (integer), RECORD_COUNT (integer), CHROM (string), POS (big integer)
- INDEX (integer)
- ALLELE (text)
- ALLELE_LENGTH (big integer)

**VARIANT_ANNOTATION**

- DATA_SET (string), ETL_LOAD_ID (integer), RECORD_COUNT (integer), CHROM (string), POS (big integer)
- RS_ID (string)
- SOURCE_TYPE (string)
- NOTES (string)

**VARIANT_FILTER**

- DATA_SET (string), ETL_LOAD_ID (integer), RECORD_COUNT (integer), CHROM (string), POS (big integer)
- FILTER_ID (string)

**VARIANT_MASTER**

- DATA_SET (string), ETL_LOAD_ID (integer), RECORD_COUNT (integer), CHROM (string), POS (big integer)
- QUAL (float)

**VARIANT_INFO**

- DATA_SET (string), ETL_LOAD_ID (integer), RECORD_COUNT (integer), CHROM (string), POS (big integer)
- *[<flexible attribute>]+*

**VARIANT_SAMPLE**

- DATA_SET (string), ETL_LOAD_ID (integer), RECORD_COUNT (integer), CHROM (string), POS (big integer)
- SAMPLE_ID (string)
- GT_INT_0 (integer)
- GT_PHASED (character)
- GT_INT_1 (integer)
- GT_REST (string)
- NONREF_ALLELE_COUNT (integer)
- *[<flexible attribute>]+*

---

**Listing 3: Content Tables**

The "VARIANT MASTER" table stores the quality "QUAL" value for all samples in this line. "VARIANT FILTER" records the filters that have been passed or failed. Each line may have several filters assigned to it, so the decision was made to normalize the data and store it in its own table instead of adding the filter information to the "VARIANT MASTER" table. Similar to filters, each line can have zero or more annotations associated with it. These are stored in the "VARIANT ANNOTATION" table. The "VARIANT ALTERNATE" table is the largest of these six tables. For each line this table stores several rows. One for each allele specified in the "REF" and "ALT" fields of a VCF file. The "INDEX" column is used to interpret, which allele is stored. The value "0" always stands for the reference allele, that is the allele that the reference genome shows at this position in the chromosome. Accordingly, the index is incremented for each alternate allele specified in the “ALT” field. The genotype information given for each sample uses the “INDEX” column to reference the allele values. Storing indices instead of the actual values for the allele spurts the advantage that researches can easily determine the share of variation in a cohort as “0” stands for the reference genome and any other value stands for a variation. The actual value of the allele is stored in the “ALLELE” column. The values stored here are any strings of arbitrary length composed of the alphabet “A”, “C”, “G”, “T”.

The two tables “VARIANT INFO” and “VARIANT SAMPLE” are flexible in structure. Listing 3 shows the base structure, which is the static part, of these tables. The “VARIANT INFO” table records all the key/value pairs specified in the “INFO” field of a line. Instead of storing key/value pairs the chosen approach utilizes the benefit of a column-oriented database to easily add column during the runtime of the system. Thus the table grows during the lifetime of the application. This approach needs an additional structure to match the dynamically created columns to the original keys from the VCF file. Therefore, a lookup table has been introduced (see Listing 4). This lookup table stores the names of the original fields in the VCF and their mapping to new created columns in the “VARIANT INFO” or “VARIANT SAMPLE” table.

**VARIANT_LOOKUP**
- DATA_SET (string), ETL_LOAD_ID (integer)
- TYPE (string)
- VCF_FIELD_NAME (string)
- INDEX (string)
- TABLE_NAME (string)
- COLUMN_NAME (string) Text

Listing 4: Lookup Table for Matching Dynamic Columns to Original Keys

Listing 5 details the pseudo-code algorithm of column matching and creation for the “VARIANT INFO” data. Similar pseudo-code is applicable to map data to the “VARIANT SAMPLE” table. The only two differences here are (1) the separators, as can be seen in the example file in Listing 1 and (2) the value strings are not in key/value format, but an extra column “FORMAT” specifies how to interpret the value strings for all the samples in the line.

```
Parse info field of VCF
Split up key/value pairs at the symbol “;”
For each key/value pair:
       Split up key/value pair into INFO_ID = key and set of values
       Lookup DATA_TYPE and NUMBER in VARIANT_INFO_MASTER for INFO_ID
       Split up set of values at “,” (use NUMBER as an indicator for max INDEX)
       If column(s) with name <INFO_ID>_<DATA_TYPE>_INDEX do(es) not exist in
        VARIANT_LOOKUP for this DATA_SET/ETL_LOAD_ID combination:
               Alter table VARIANT_INFO add column(s) <INFO_ID>_<DATA_TYPE>_INDEX
               Update VARIANT_LOOKUP table with according information
       Insert data into VARIANT INFO table
```

Listing 5: Pseudo-code for Column Matching and Creation
