## Supplementary material for "The Next Generation Precision Medical Record - A Framework for Integrating Genomes and Wearable Sensors with Medical Records": s1-vcf_specification

As market leader in enterprise application software, SAP SE (NYSE: SAP) helps companies of all sizes and industries run better. From back office to boardroom, warehouse to storefront, desktop to mobile device – SAP empowers people and organizations to work together more efficiently and use insights to effectively deliver tangible results or stay ahead of the competition. SAP applications and services enable more than 282,000 customers worldwide to operate profitably, adapt continuously and grow sustainably. As a leader in cloud technology, SAP follows privacy and security standards to the highest degree. SAP solutions and technologies help healthcare organizations engage better, collaborate better, and decide better – to deliver cost-effective care with the best outcomes so that people can live more healthily.

SAP's latest technology, SAP HANA®, is a flexible, multi-purpose in-memory data management and application platform that has completely transformed the relational database industry. By providing advanced capabilities, such as text analytics, spatial processing and data virtualization on the same architecture, it further simplifies application development and processing across big-data sources and structures. This makes SAP HANA a highly suitable platform for building and deploying next-generation, real-time applications and analytics. SAP HANA has already been adopted by global health and medical organizations (and thousands of other institutions) worldwide. With office locations in more than 130 countries, focused teams are working around the clock to develop solutions that enable Precision Medicine.

The SAP Foundation for Health is built on top of the SAP HANA platform. The SAP Foundation for Health provides a flexible and extensible clinical data warehouse model, industry aware data integration management, and real-time analytics on large-scale structured and unstructured data. Unlike traditional data warehouses, the Foundation for Health comes with built-in services to facilitate better diagnosis and decision support. It provides an open and secure platform for genomics, patient cohort building, patient trial matching and further care collaboration solutions developed by SAP and partners. The SAP Foundation for Health leverages generic SAP HANA tools and components, such as SAP Data Services, an Extract, Transform & Load (ETL) tool that can be applied to load all kinds of health data. Thanks to the generic healthcare data model, data from many different sources including 3rd party vendor databases, clinical information systems, tumor registries, biobank systems and even unstructured text documents can be integrated. This data can be processed by applications developed on top of the foundation.

One of the first SAP applications running on top of this platform is the award winning SAP Medical Research Insights. With Medical Research Insights, clinical researchers can filter and group patients according to certain attributes, select patients for clinical trials based on certain criteria, perform Kaplan-Meier estimations on the fly and view individual patient timelines in a graphical way. Medical Research Insights was built in cooperation with Nationales Centrum für Tumorerkrankungen (NCT) in Heidelberg, Germany. In October 2013, SAP received special recognition from the White House Office of Science and Technology Policy for SAP's contribution with the Stanford School of Medicine and NCT in accelerating the promise of personalized medicine. More recently, the American Society of Clinical Oncology selected SAP as the technology partner to develop CancerLinQ™, a cloud-based big data platform. CancerLinQ will unlock real-world patient care data from millions of electronic health records and more securely process and analyze the data to provide immediate high-quality feedback and clinical decision support to providers.

An integral component of Foundation for Health is the genome data schema, which stores all data from the Variant Call Format and standard annotation formats. By storing all the variant and associated metadata in a set of tables, the data can be easily queried or integrated with additional annotations in a scalable fashion. This novel approach has been tested using the phase 3 release of the 1000 Genomes Project dataset (2504 whole human genomes) and enables real-time queries across all variant data. An extensible UI framework provides an intuitive way to visualize and explore genome data in the context of the reference genome and annotation data.

With wearable technology becoming more widespread, digital services can promote remote patient care, behavioral monitoring and potentially enable preventative medicine. SAP Health Engagement enables preventative and remote care for chronic diseases and conditions. Recently, SAP partnered with Roche Diagnostics GmbH and developed SAP Health Link – a solution that enables health care providers to connect with Type II Diabetes (T2D) patients through digital services. From a web-based portal, doctors are able to onboard and set goals for their patients. Using a mobile app, patients can monitor and record health parameters, either actively or by passively connecting to wearable's and sensors. The app helps patients adhere to required medications, and patients can order medication directly from the app. Doctors can remotely monitor patients' health parameters and communicate with patients directly if any outliers or unusual patterns are observed. This solution can enable early detection of patients at risk and promote lifestyle and behavioral changes in patients. Researchers can use the analytical dashboards to study the effectiveness of health programs. SAP Health Link, now live at Roche, is a Health Engagement solution built on the SAP HANA Cloud Platform. Although the app is tailored to T2D, the SAP Health Engagement flexible development toolkit with can be used to develop similar apps focusing on other chronic diseases and conditions.

SAP's collaboration with Stanford School of Medicine aims to leverage SAP's technologies to achieve a better understanding of global human genome variation and its implications in disease, particularly cardiovascular disease. Led by Carlos D. Bustamante, professor and director of the Stanford Center for Computational, Human and Evolutionary Genomics and Euan A. Ashley, associate professor of Medicine and Genetics, the collaboration aims to advance genomics research in the clinical environment, ultimately leading to improved healthcare and personalized medicine. More recently, SAP worked with Stanford's Clinical Genomics Service to integrate genomic and clinical data, with the goal of enabling actionable insights that support more precise, personalized clinical care. Using SAP HANA, Stanford researchers were able to integrate genomic, clinical, and wearable data of a select number of patients. By integrating these datasets and automating the filtering and prioritization of genetic variants, researchers could quickly identify clinically actionable mutations, optimal drug dosages, and potentially new mutations playing a role in a patient's condition.
